# Exploring the Application of the MoCMC System for Determining In Vitro Product Comparability

**DOI:** 10.64898/2025.12.10.693525

**Authors:** Marilyn N. Martinez, David G. Longstaff

## Abstract

**Purpose:** To explore the utility of the Matrix of Chemistry, Manufacturing and Control (MoCMC) approach for evaluating product physicochemical (Q3) similarity as part of a product bioequivalence assessment.

**Methodology:** Each Q3 parameter was converted to a radius of a polygon using a mathematical equation that converted each reference Q3 parameter to a value of “5”. That equation was applied to the corresponding test product Q3 parameter and the areas of the treatment polygons determined. The ratio of the product area/outer area [Matrix Comparability Index (MCI)] was calculated using an outer polygon with each radius = “10”. When generated across multiple lots for each treatment, the MCI values were statistically compared. Both actual (from our previously published investigation) and hypothetical MCI values were used to characterize the performance of the MoCMC across a range of situations.

**Results:** MCI values successfully captured product Q3 differences, with the influence of any Q3 parameter decreasing as the number of radii increases. The ability to identify statistically significant product differences decreased as variability of either test or reference MCI values increased. By combining statistical analysis with a comparison of the spread of test and reference MCI values, the MoCMC provided a sensitive tool for assessing product comparability and identified product differences previously missed by comparing *in vitro* dissolution profiles alone

**Conclusions:** While the impetus for this research was to identify a tool for supporting the evaluation of product bioequivalence containing locally acting drugs, its strengths support its potential use across a wider scope of situations.

## Introduction

The bioequivalence (BE) assessment of locally acting, non-systemically absorbed, active pharmaceutical ingredients (API’s) presents the regulatory challenge of systemic drug concentrations that do not reflect the relative drug concentrations at the site of action. This situation has led to a reliance on terminal clinical endpoint BE studies for veterinary antiparasitic drug products whose therapeutic target resides within the lumen of the gastrointestinal (GI) tract.

The Food and Drug Administration Center for Veterinary Medicine (FDA CVM) recently conducted a study to explore the development of a scientifically robust alternative to clinical endpoint BE studies^1^. That research explored the question of whether *in vitro* dissolution could serve as a surrogate for comparing product relative rates and extent of *in vivo* tablet dissolution within the canine GI tract. To avoid the need to use clinical endpoints, the study examined the relative bioavailability of three experimental monolithic tablet formulations containing the two poorly soluble, highly permeable APIs, ivermectin (IVM) and praziquantel (PRZ). The assumption for both compounds was that since *in vivo* dissolution was the rate limiting step in their oral absorption, *in vitro* dissolution could be used to predict *in vivo* dissolution and therefore tablet formulation effects on PRZ and IVM bioavailability. Based upon that assumption, the rates and extent of *in vivo* drug exposure was used as a biomarker for comparing the rates and extent of *in vivo* dissolution across the three experimental formulations.

A 3-period, 3-treatment, 3-sequence crossover study was conducted in 27 healthy Beagle dogs. Pairwise treatment comparisons of the rates and extent of systemic drug concentrations were conducted for each of the two active ingredients (IVM and PRZ). The experimental tablets used in that investigation were developed to exhibit the following distinct release characteristics^2^:

Treatment A (Trt A): Fast release of both IVM and PRZ

Treatment B (Trt B): Fast release PRZ and medium release IVM

Treatment C (Trt C): Medium release PRZ and fast release IVM

To manufacture these three treatments, each API was individually granulated with excipients. The same formulation was used for the respective API fast and medium release granules. Granule size was intentionally varied to further adjust PRZ or IVM *in vitro* release rates. The granules for both drugs were combined and compressed to form each of the three treatments. While ideally, the reference and generic formulations would have the same active and inactive ingredients (Q1) in similar proportions (Q2) and exhibit the same physicochemical characteristics (Q3), the deviations from Q1, Q2 and Q3 comparability was necessary to ensure inequivalence for some between-product comparisons. This inequivalence provided an opportunity to assess whether *in vitro* dissolution data, when generated across a range of conditions, could identify *in vivo* between-product differences.

The *in vitro* dissolution profiles of the three treatments were compared in media consisting of 0.1N HCl + 0.2% sodium lauryl sulfate (SLS), pH 4.5 acetate buffer + 0.5% SLS, and 0.01M pH 6.8 phosphate buffer + 0.5% SLS. Except for the test results in 0.1 N HCL (where the % IVM dissolved from Trts A and C increased and subsequently decreased), Trt A exhibited the same or faster IVM and PRZ *in vitro* release as compared to that of the other two formulations.

Based upon the results of the in vitro dissolution studies, the expectation was the PRZ component of Trt A would be bioequivalent (BE) to the PRZ component of Trt B and that both Trts A and B would have higher peak exposure (and potentially higher total drug exposure) as compared to that of Trt C. With regard to IVM, the expectation was that Trt A would be BE to Trt C and that both would have higher peak exposure and similar or greater total IVM exposure as compared to that of Trt B. Although the *in vitro* dissolution results for the individual tablet components were predictive of PRZ and IVM relative total drug exposure between Trts B vs C, it failed to predict the much lower relative bioavailability (rate and extent) of Trt A^1^. The IVM comparative in vitro dissolution profiles were also inconsistent with the slightly greater peak exposure of Trt B vs C. These in vivo/in vitro inconsistencies highlighted the error that could occur when relying solely on *in vitro* dissolution for assessing product comparability. The physiologically based pharmacokinetic modeling simulation software GastroPlus (v9.9) was used to explore potential reasons for this outcome (Martinez et al., submitted)^3^. While that modeling effort enabled us to exclude certain possibilities for the observed *in vivo* / *in vitro* discrepancies, numerous unanswered questions remained.

The Q3 parameters included in a BE product assessment should reflect the product’s critical quality attributes (CQAs). However, an assessment of product sameness can be difficult when considering multiple Q3 parameters or when there is inherent variability across lots of the reference product. To addressing this challenge, we explored the applicability of the Matrix of Chemistry, Manufacturing and Control (MoCMC).

The MoCMC system is derived from the Sediment Delivery Model (SeDeM) system which was introduced in approximately 2005 as a mechanism for supporting formulation development^4^ and has been used to support the optimization of oral product formulations.^5,6,7,8,9^ Both systems provide a visual and mathematical platform for simultaneous comparing multiple attributes across multiple formulations. Used by scientists at Texas A&M University during a collaborative research effort with CVM^10^, the MoCMC system provided the basis for identifying inequivalent experimental formulations of products intended for intramammary infusion (a locally acting dosage form indicated for the treatment of bovine mastitis).^11,12^

## Methods

The MoCMC and SeDeM approaches involve the mathematical conversion of Q3 parameter values into radii originating from the center of a polygon. For the reference product, the radius for each Q3 parameter is converted to a value of “5”^8^ via mathematical equations. Each reference-derived equation is used to convert the corresponding test product Q3 parameters into radii. The sets of radii for each lot of the test or reference products define the shape of a polygon whose internal area can be measured. That internal polygonal area represents the product fingerprint. The product fingerprint is divided by an outer polygonal area where the radii of the same Q3 parameters are given a value of “10”. The ratio of the product fingerprint area/outer surface area is termed the Matrix Comparability Index (MCI).

### Determining the Radius for Each Attribute Used in the Comparison of Trts A, B and C

Our assessments were limited by the Q3 information available for Trt A, B and C. Therefore, although hardness and friability are typically of interest, those measurements were not available for inclusion in this MoCMC assessment. This resulted in a preponderance of radii defined by dissolution results. Despite this deficiency, it was considered of value to determine if the resulting treatment MCI comparisons would have been consistent with the aforementioned outcomes of the *in vivo* BE study.

Six radii were included in the MoCMC assessments: tablet dissolution in 2 sets of media; at 20 and 120 min in 0.1N HCl + 0.2% SLS and acetate buffer pH 4.6 + 0.5% SLS, granule diameter, and the % theoretical concentration corresponding to the various sieve cuts. For IVM, to accommodate the increase and subsequent decrease in Trts A and C % dissolved in 0.1N HCL + 0.2% SLS, the % dissolved at 120 min was divided by the amount dissolved in 20 min. When considering the elimination of one set of dissolution characteristics (e.g., % dissolved at 120 minutes), the MoCMC would have been based on only 4 parameters which, for reasons identified in the Results, was considered too few to provide a reliable outcome.

The equation used to describe the reference radius for each parameter is provided in Supplemental Table 1. Unless otherwise specified, the reference formulation was designated as Trt A and the test formulations as Trt B or C. Data for IVM and PRZ were evaluated separately.

### Determining Polygonal Areas

Using the calculated treatment radii, a polygonal plot was generated for each treatment within a single figure using the radar plot function in Excel (Microsoft^®^ Excel^®^ for Microsoft 365 Version 2502). The plot was saved in JPEG format and imported into ImageJ (ImageJ 1.54g, National Institutes of Health, USA, http://imagej.org). The lines defining the sides of each polygon were re-traced using the polygonal tool within ImageJ to enable polygonal area calculation without scaling (i.e., since all plots were contained within the same figure, the relationship between radius length and pixel distance would be identical).

### Method Characterization

To ascertain the utility of the MoCMC system for supporting the BE of products containing non-systemically absorbed drugs, it was necessary to explore: 1) the influence of the number of parameters included in the assessment; 2) the discriminative capability of the treatment comparison as a function of the magnitude of product MCI differences versus the test vs reference MCI variability.

#### Sensitivity to the number of attributes

To assess the influence of the number of parameters included in the MoCMC on the sensitivity to parameter differences (when expressed as the ratio of treatment mean MCI values), we generated MCI values when only parameter (radius) had a disparate value in sets where the total number of parameters (radii) ranged from 4 – 6.

#### Influence of treatment MCI variability and differences between means

To explore factors that could affect the outcome of the MoCMC-based comparisons, 1000 normally distributed test and reference MCI values were generated via Monte Carlo simulation [Oracle Crystal Ball software (release 11.1.3.0.000)]. Specific details of each challenge are provided in the Results.

From these 1000 simulated test and reference values, five sets of test and reference values were randomly selected. Except as noted in the Results, each set contained 5 reference “lots” and 3 test “lots”, with each “lot” having its own MCI value. Limiting the number of test and reference “lots” to n = 3 and n = 5 per set, respectively, is intended to reflect the number of reference and pilot test lots typically available to support veterinary product BE applications. Given the small number of MCI values included per set, each set of randomly selected values was observed to differ from the population means and variability specified for generating the Monte Carlo simulations. Therefore, within each set, the distribution of individual values, product means and their corresponding % coefficient of variation (%CV) was determined to better understand the results of the statistical analysis.

The following simulated populations were used to explore the influence of variability on the ability to detect inter-product differences:

1) Hypothetical populations: Test and reference mean MCI values differed by 10%. A value of 10% was selected because as the % difference increases, the ability to detect differences would likewise increase. Comparisons included the following:
  a. Reference (Ref) CV = 5% and:
    i. The test CV was 5%.
    ii. The test CV was 20%
  b. The Ref CV was 10% and:
    i. The test CV was 5%.
    ii. The test CV was 20%
2) Exploring the formulations used in the IVM/PRZ *in vivo* study:
  a. Potential for declaring IVM or PRZ Trt C equivalent to Trt A: 5 sets of randomly selected treatment values were simulated. Both for PRZ and IVM, comparisons were limited to Trts A vs C because for both APIs, the difference between Trts A and C were less than that between Trts A vs B. Since all treatments failed to demonstrate *in vivo* BE to Trt A, selecting the comparison with the smallest difference provided the greatest challenge to our ability to identify potential limitations in the MoCMC system. For each set of simulations, the population mean values were defined by the MCI value obtained from the single lot of the actual treatments. The population %CV was set as: 1) 20% or 10% for Trt A and 10% for Trt C (PRZ) and 2) 20% for Trt A and 1% or 10% for Trt C (IVM).
  b. Potential for declaring PRZ or IVM Trts B equivalent to Trt C: 5 sets of randomly selected treatment values were simulated. In this comparison, Trt B was arbitrarily selected to be the reference product. The simulated population mean values were defined by the MCI value obtained with the single lot of the actual treatments. The population %CV was simulated as 10% for Trt B and 1% or 10% for Trt C (PRZ) and 20% for Trt B and 1% or 10% for Trt C (IVM). Because of the magnitude of difference in the mean MCI values for IVM Trts B and C, to further challenge this evaluation, the %CV for Trt B was 20% and the %CV for Trt C was 1% or 10%.

The use of different simulation %CV input values supported an examination of relationships between differences in treatment MCI versus their respective variabilities on comparability conclusions generated with an MoCMC analysis.

#### Statistical analysis

Statistical analysis was conducted in SAS v 9.4 using Proc mixed (class treatment; model MCI=trt; repeated/group=trt; LSMeans trt/diffs Cl alpha = 0.01). Each set was analyzed separately. In addition, each pairwise comparison was generated separately. We note that this is a statistical test for differences and not of equivalence.

## Results

### Method Characterization

#### Sensitivity to the number of attributes

An increase in the number of attributes included in the polygonal area reduced the magnitude of influence exerted by a single parameter on the MCI value (Table 1, Figure 1a,b). For Hypothetical Test 1 (HT-1), since its MCI value was less than that of Hypothetical Reference (H-R), as the number of parameters increased, the MCI ratio of H-R/HT-1 dropped from over 110% to 107%. For Hypothetical Test 2 (HT-2) and Hypothetical Test 3 (HT-3) whose MCI values were greater than that of H-R, as the number of parameters increased, their ratios relative to H-R increased closer to 100%. Despite the trend for more parameters reducing the impact of a single discordant parameter, little further change in impact was observed when the number of radii increased from 5 to 6.

**Figure 1:**
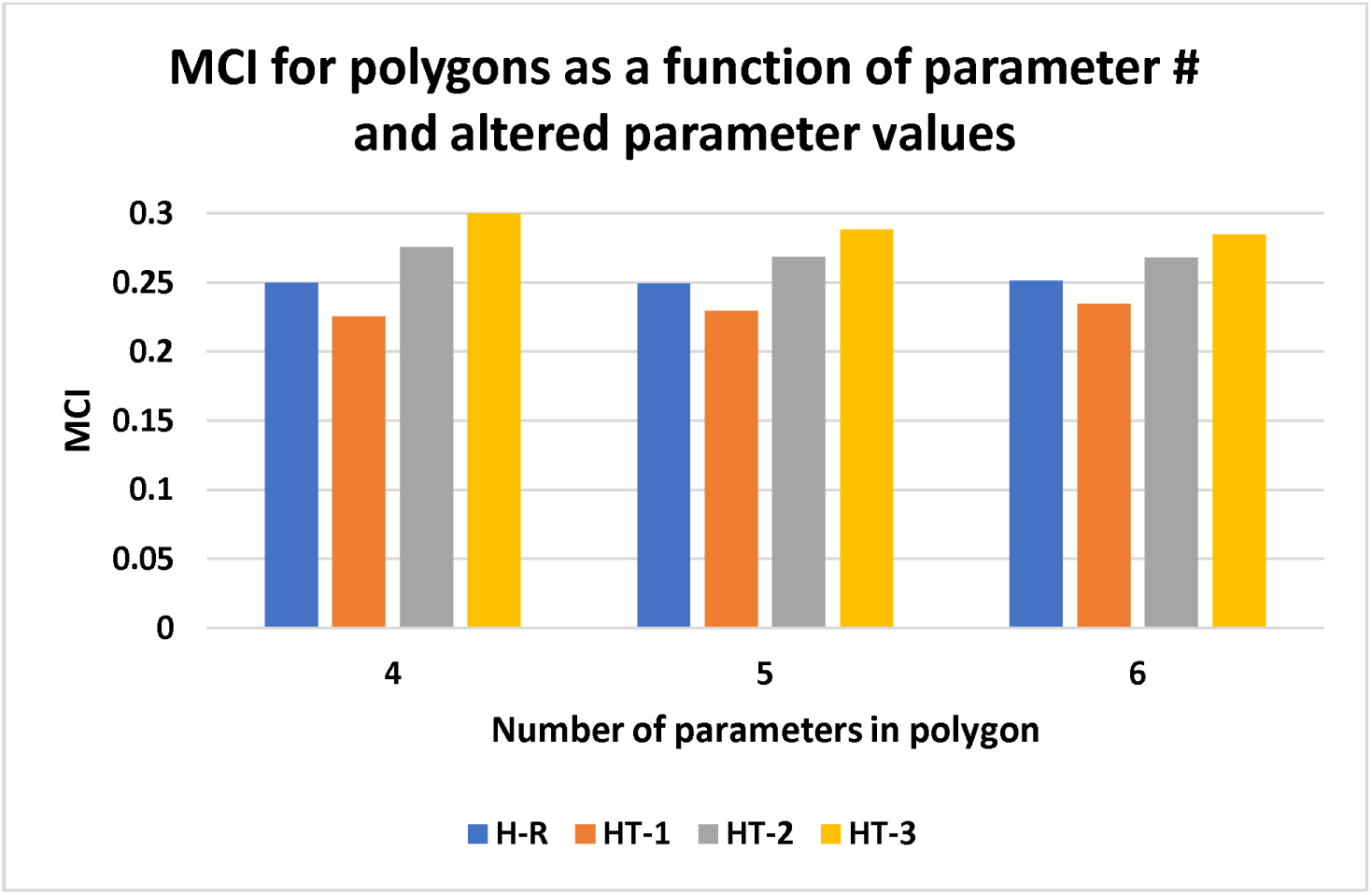
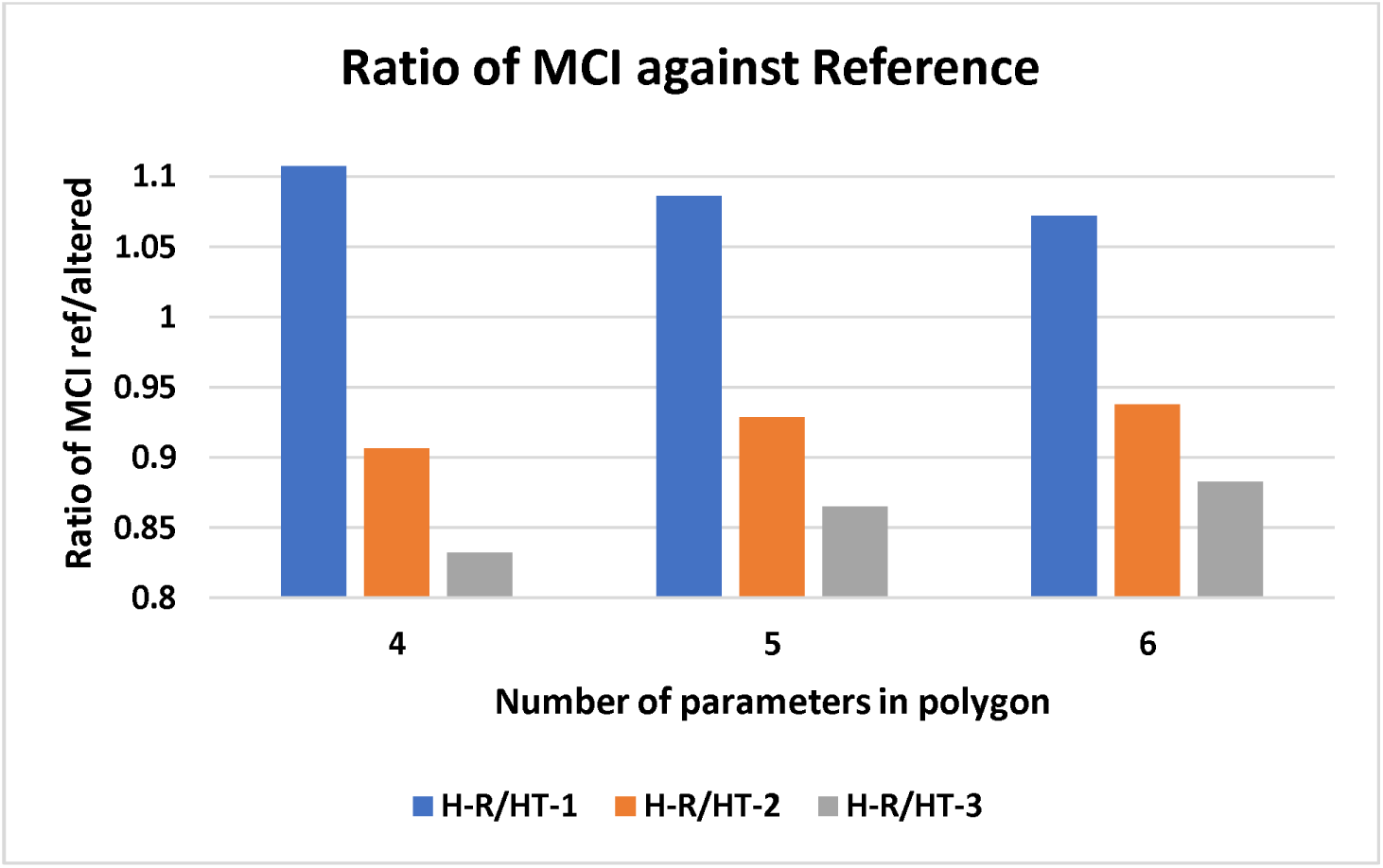
Influence of the number of parameters on the comparison of MCI values: a) MCI for polygons as a function of number of parameters and altered parameter values, b) treatment MCI values expressed relative to the reference.

**Table 1:**
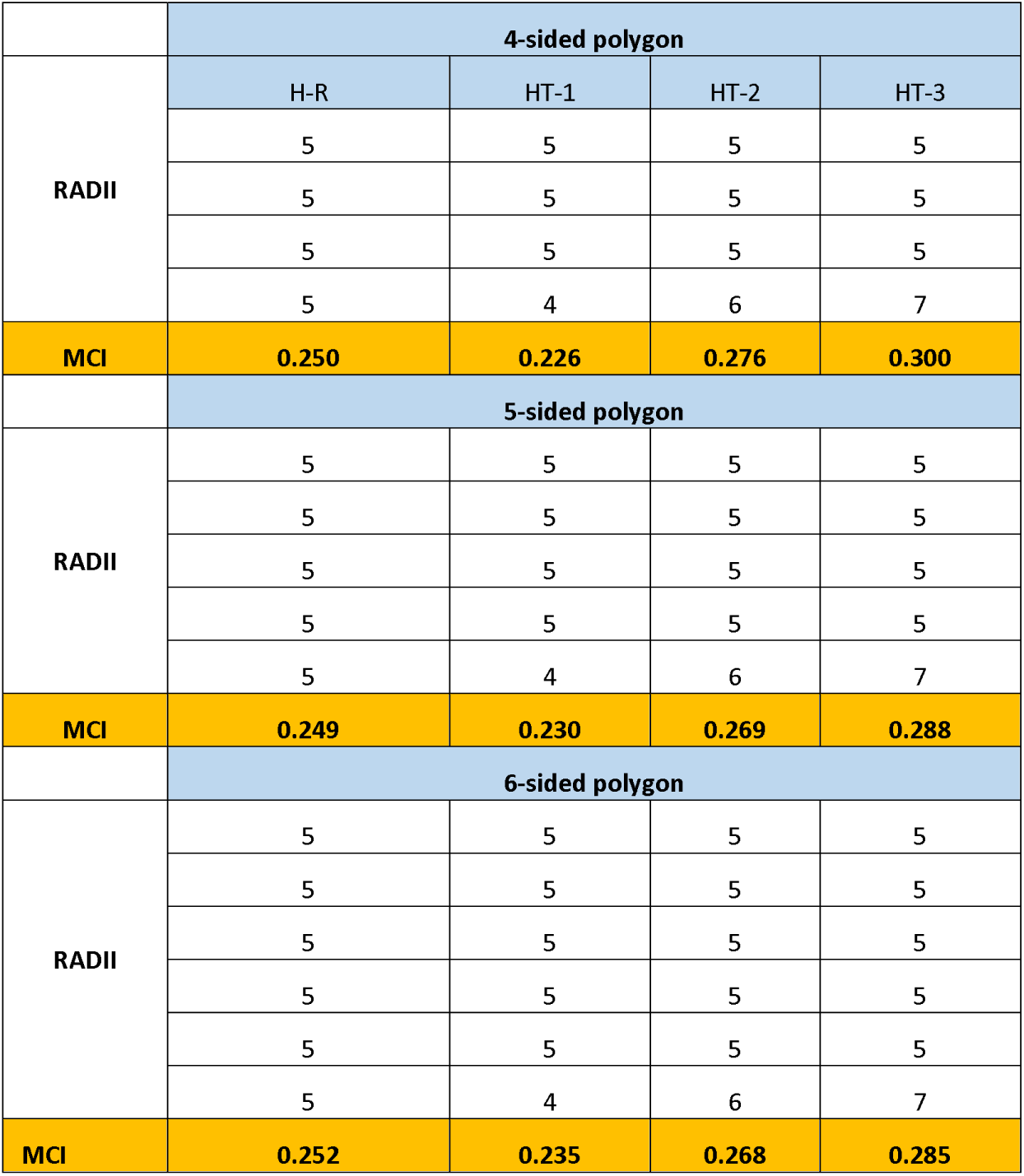
Influence of the number of radii on the MCI value (slight deviations from H-R = 0.250 was attributable to small errors associated with areas calculated using ImageJ).

### Influence of Parameter Values on MCI Estimates and on Statistical Assessments

#### Influence of treatment MCI variability and differences between means

All simulations included a 10% difference between population Test and Ref MCI values.

Although the simulations were conducted with a test product %CV up to 20%, limiting the random selection to n = 3 lots per set resulted in only three sets approaching the targeted level of variability or ratios approaching the targeted difference in product means. The mean and %CV for each set is provided in Table 2a.

**Table 2a:**
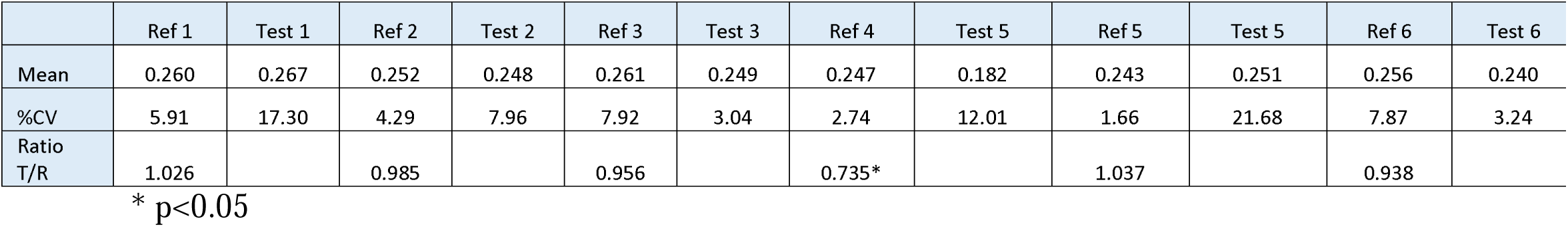
Test and reference MCI comparisons: Sets 1 – 5 were selected from the simulation where Ref %CV was set at 5% and Test %CV at 20%. Set 6 was the one set where statistical significance was not observed when both the Test and Ref products were simulated based on a population CV of 5%.

|  | Ref 1 | Test 1 | Ref 2 | Test 2 | Ref 3 | Test 3 | Ref 4 | Test 5 | Ref 5 | Test 5 | Ref 6 | Test 6 |
| --- | --- | --- | --- | --- | --- | --- | --- | --- | --- | --- | --- | --- |
| Mean | 0.260 | 0.267 | 0.252 | 0.248 | 0.261 | 0.249 | 0.247 | 0.182 | 0.243 | 0.251 | 0.256 | 0.240 |
| %CV | 5.91 | 17.30 | 4.29 | 7.96 | 7.92 | 3.04 | 2.74 | 12.01 | 1.66 | 21.68 | 7.87 | 3.24 |
| Ratio<br>T/R | 1.026 |  | 0.985 |  | 0.956 |  | 0.735* |  | 1.037 |  | 0.938 |  |

\* $p < 0.05$

**Table 2b:** Test and Ref MCI comparisons: Sets 1 – 5 were selected from the simulation where Ref %CV was set at 10% and Test %CV at 5%.

|  | Ref 1 | Test 1 | Ref 2 | Test 2 | Ref 3 | Test 3 | Ref 4 | Test 4 | Ref 5 | Test 5 |
| --- | --- | --- | --- | --- | --- | --- | --- | --- | --- | --- |
| Mean | 0.254 | 0.221 | 0.226 | 0.225 | 0.256 | 0.240 | 0.242 | 0.221 | 0.238 | 0.221 |
| %CV | 9.97 | 4.49 | 13.10 | 2.22 | 7.87 | 3.24 | 10.13 | 2.87 | 6.30 | 1.65 |
| Ratio<br>T/R | 0.868* |  | 0.992 |  | 0.938 |  | 0.913* |  | 0.928 |  |

\* $p < 0.05$

**Table 2c:** Test and Ref MCI comparisons: Sets 1 – 5 were selected from the simulation where Ref %CV was set at 10% and Test %CV at 20%.

|  | Ref 1 | Test 1 | Ref 2 | Test 2 | Ref 3 | Test 3 | Ref 4 | Test 4 | Ref 5 | Test 5 |
| --- | --- | --- | --- | --- | --- | --- | --- | --- | --- | --- |
| Mean | 0.254 | 0.267 | 0.226 | 0.248 | 0.256 | 0.249 | 0.242 | 0.182 | 0.238 | 0.251 |
| %CV | 9.97 | 17.30 | 13.10 | 7.96 | 7.87 | 3.04 | 10.13 | 12.01 | 6.30 | 21.68 |
| Ratio<br>T/R | 1.050 |  | 1.097 |  | 0.974 |  | 0.752* |  | 1.057 |  |

\* $p < 0.05$

When Test and Ref populations were simulated at %CV = 5, statistically significant differences were observed across all but one set of comparisons (Ref 6 and Test 6 in Table 2a). When the Ref was simulated with a %CV=10 but the Test simulation was maintained with a 5% CV, the number of sets for which significance was detected decreased from 4/5 to 2/5 (Sets 1 and 4). Keeping the Ref at a 10% CV but increasing the Test to 20% CV further reduced the ability to detect statistically significant differences (Table 2c). Therefore, regardless of which product exhibits the higher %CV, an increase in the between-lot variability reduces the power to identify statistically significant treatment differences.

Given the potential for inadequate statistical power to detect treatment differences due to the limited number of treatment observations and to the magnitude of variability associated with the test and/or reference products, it was necessary to consider the utility of employing an alternative (non-statistical) approach for generating the between-treatment comparison. That approach entailed plotting the individual “lot” values of the Test and Ref formulations and determining if the MCI values for the Test formulation were contained within the range of Reference MCI values. Using that approach, only Sets 3 and 6 had all test lot MCI values contained within that of the Ref. It is also noteworthy that in those two cases, the %CV of the Test product was less than that of the Ref, suggesting that the 6-7% differences in product MCI values will be more difficult to detect (visually or statistically) when the Test product MCI %CV is less than that of the Ref (Figure 2a).

**Figure 2a:**
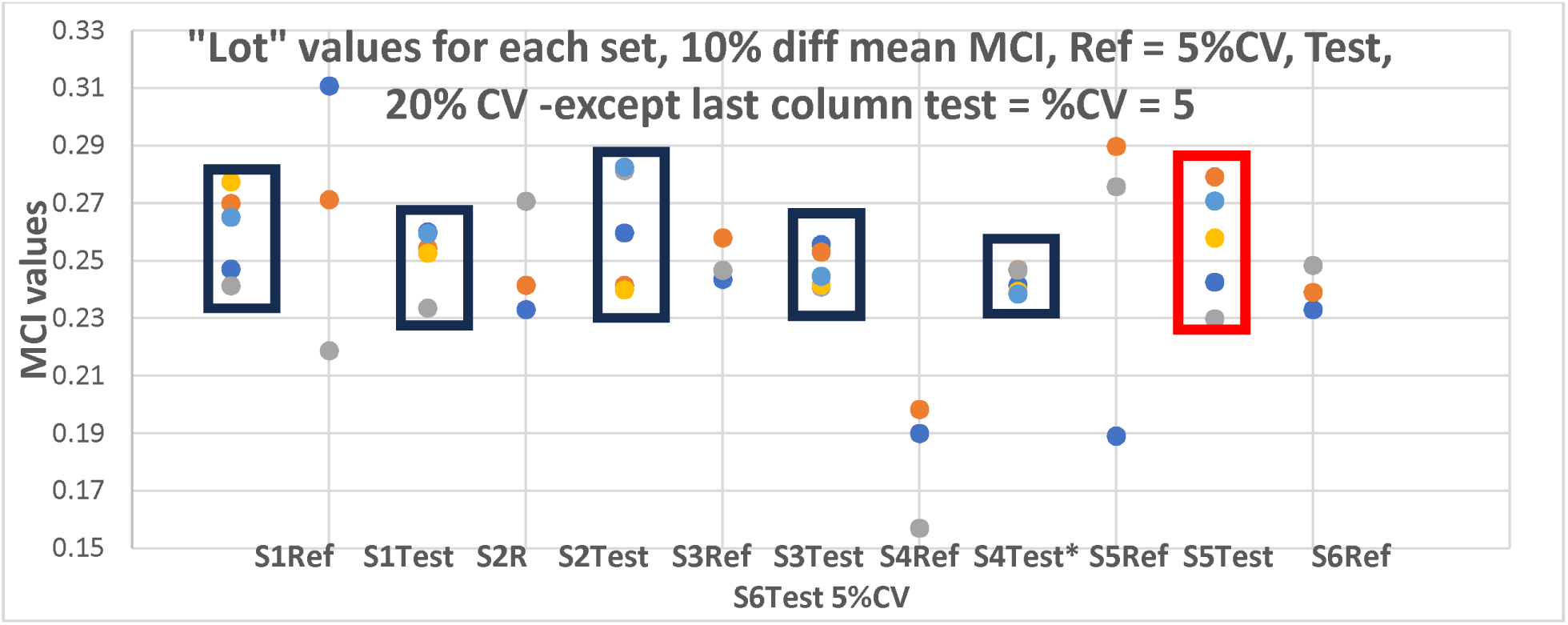
when Test and Ref products were simulated with a 10% difference in mean MCI values, Ref = 5%CV (n=5 per set), Test, 20% CV (n=3 per set), except for the last column where the test CV= 5% (n=3 per set. Sets highlighted with black boxes represent the Ref product. The Ref set derived from the prior Monte Carlo simulation, representing the one set of comparisons for which statistically significant differences were not detected (S6Ref) is highlighted with the red box (n=5). The corresponding test values is titled S6Test 5%CV (n=3).

When the Ref product CV was increased to 10%, statistically significant differences were typically not observed (Figures 2b and C). Note that to focus on the influence of comparative Test versus Ref product variability, the same five randomly selected reference lots were used in each of these evaluations within a given %CV.

Although generating thousands of bootstrap sets within each simulated condition would have described the potential distribution of outcomes, outcome distribution was not the objective of this assessment. Rather, the limited number of sets included in this analysis enabled an assessment of how the spread of “lots” within each set might influence conclusions derived from the MoCMC approach in the presence of limitations on “n” per treatment.

**Figure 2b:**
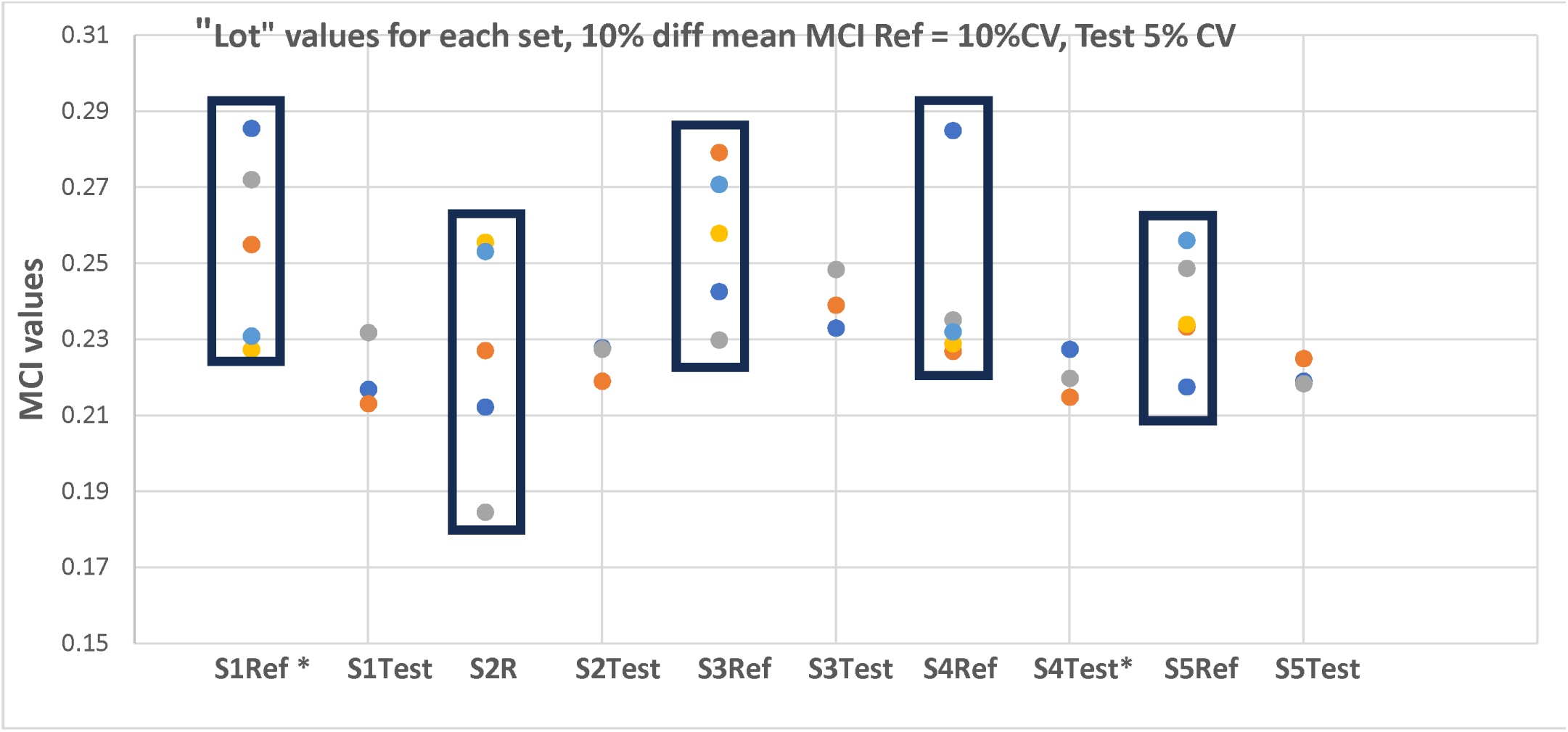
when Test and Ref products were simulated with a 10% difference in mean MCI values, Ref = 10%CV (n=5 per set), Test 5% CV (n=3 per set). Sets with a black box represents the Ref product.

**Figure 2c:**
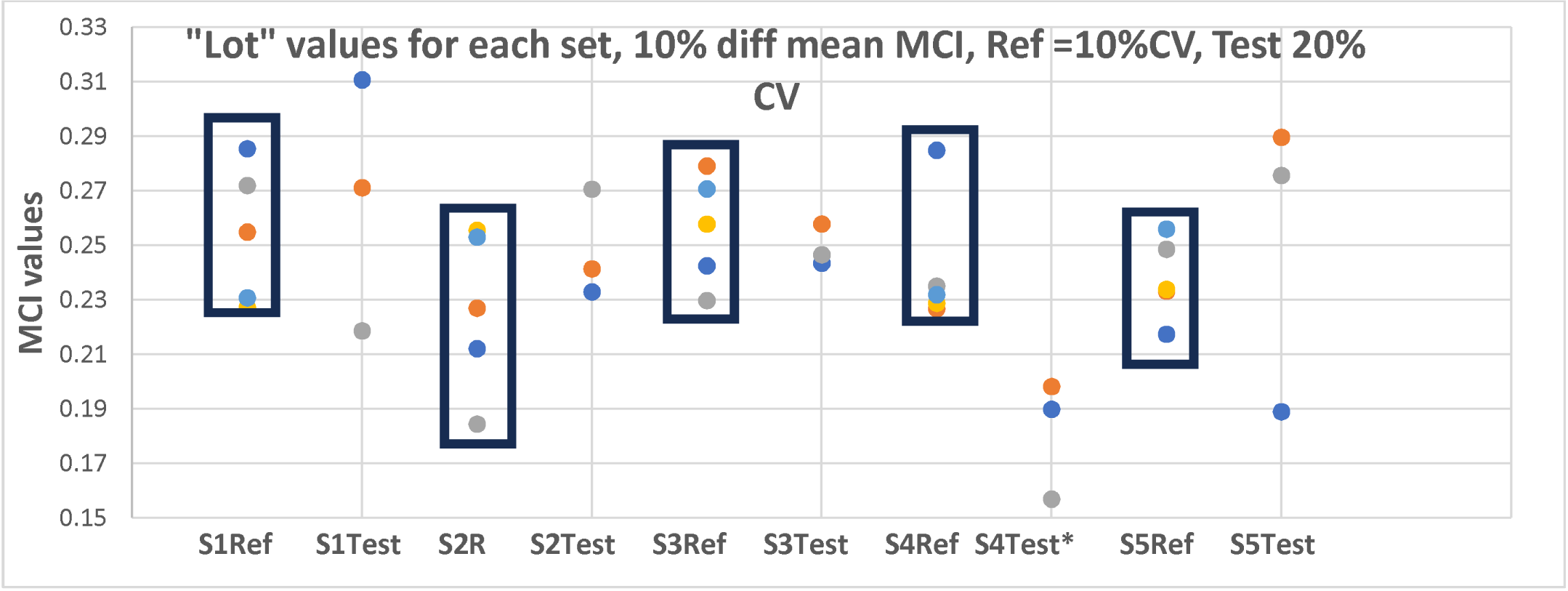
“Lot” values for each set when Test and Ref products were simulated with a 10% difference in mean MCI values, Ref =10%CV (n=5 per set), Test 20% CV (n=3 per set). Sets with a black box represents the Ref product.

Figures 2a-c demonstrated that a focus on statistical significance alone could be misleading. In particular, there were instances where significance was not achieved but the distribution of Test lots within each set differed from that of the Ref. Of concern was that despite similar mean values, there were instances where the Test product had lots whose MCI values were outside the bound defining the corresponding Ref. Accordingly, we concluded that when using the MoCMC, it was necessary to include two tiers for any pairwise comparison: 1) pairwise T-test; and 2) visual determination of whether the Test lots all are contained within the range of the Ref lots. If that two tiered process was employed, sets 1, 2, 4 and 5 would have been declared different in Figure 2a, sets 1 and 4 would have been declared different in Figure 2b, sets 1, 2, 4 and 5 would have been declared different in Figure 2c.

### Determining the Radii for Each Attribute Used in the Comparison of Trts A, B and C

#### MCI for PRZ

The radii associated with the 6 parameters included in this assessment, and the corresponding MCI values for PRZ, are provided in Tables 3a and b. Due to the lack of availability of multiple lots for any of these treatments, it was not possible to generate pairwise statistical comparisons in the absence of additional Monte Carlo simulations based upon assumptions pertaining to variability in MCI values (shown later).

**Table 3a:** PRZ parameter radii for Treatments A, B and C.

|  | % Dissolved at 20 min 0.1 N HCl | % Dissolved at 20 min acetate | Granule size | % theoretical concentration granule | Dissolved 120 min 0.1N HCl | Dissolved 120 min acetate |
| --- | --- | --- | --- | --- | --- | --- |
| A | 5.00 | 5.00 | 5.00 | 5.00 | 5.00 | 5.00 |
| B | 5.14 | 4.76 | 9.4 | 5.39 | 5.41 | 4.94 |
| C | 3.68 | 2.29 | 13.73 | 5.86 | 4.92 | 4.61 |

**Table 3b:** PRZ parameter radii for Trts A, B and C.

| Trt | MCI | Ratio to A |
| --- | --- | --- |
| A | 0.25 | 1.00 |
| B | 0.33 | 1.34 |
| C | 0.31 | 1.26 |

The polygons associated with Treatments A, B and C are shown in Figure 3.

**Figure 3:**
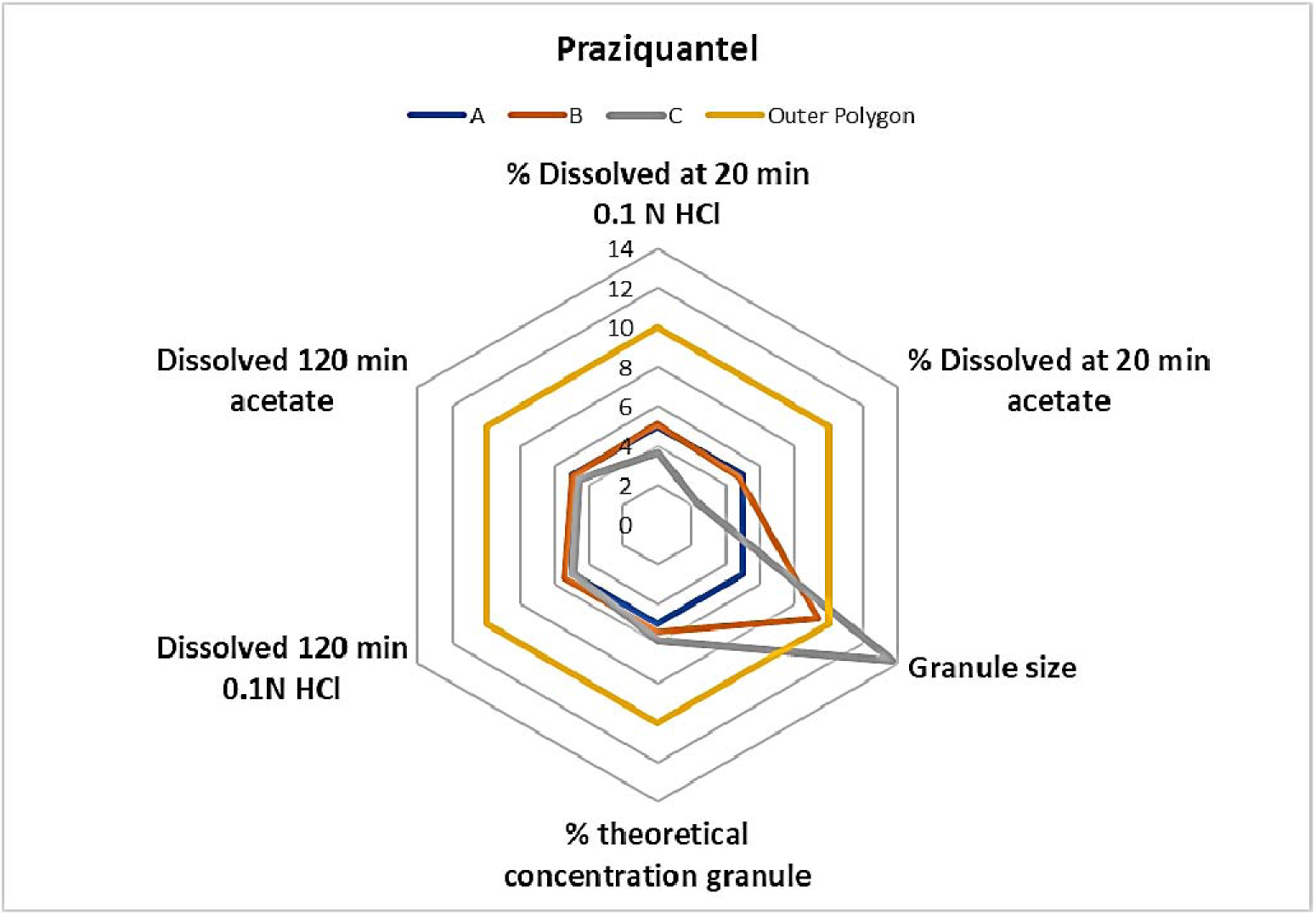
PRZ polygons for Trts A, B and C.

#### MCI for IVM

The polygons associated with Trts A, B and C are shown in Figure 4.

**Figure 4:**
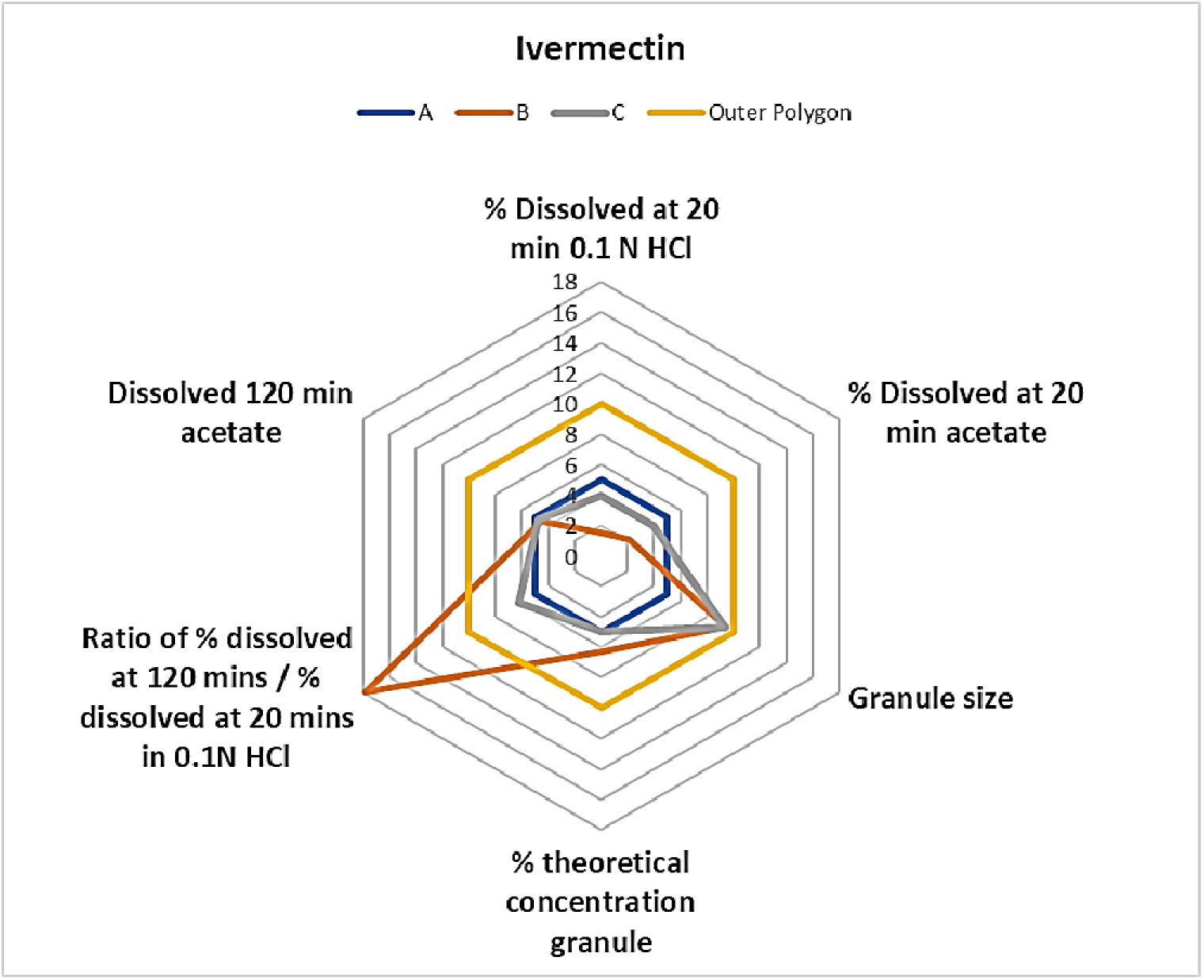
IVM polygons for Trts A, B and C.

The radii for the 6 parameters included in this assessment, and the corresponding MCI values for IVM, are provided in Table 4a and b. As with PRZ, lack of availability of multiple lots prohibited pairwise statistical comparison of three formulations.

**Table 4a:** IVM parameter radii for Treatments A, B and C.

| Trt | % Dissolved at 20 min 0.1 N HCl | % Dissolved at 20 min acetate | Granule size | % theoretical concentration granule | Ratio of % dissolved at 120 min/ % dissolved 20 min | % Dissolved 120 min acetate |
| --- | --- | --- | --- | --- | --- | --- |
| A | 5.00 | 5.00 | 5.00 | 5.00 | 5.00 | 5.00 |
| B | 1.52 | 2.10 | 9.40 | 6.25 | 17.88 | 4.54 |
| C | 3.91 | 3.98 | 9.4 | 5.00 | 6.23 | 4.73 |

**Table 4b:** IVM parameter radii for Treatments A, B and C.

| Trt | MCI | Ratio to A |
| --- | --- | --- |
| A | 0.25 | 1.00 |
| B | 0.47 | 1.92 |
| C | 0.30 | 1.22 |

### Exploring the limitations associated with the MoCMC

#### Challenge 1: Potential for declaring IVM or PRZ Trt B or C equivalent to Trt A

Monte Carlo simulations were performed using population means equaling the calculated treatment MCI values from the original study batches [Tables 3b PRZ Trt A = 0.25, Trt C = 0.31) and 4b (IVM Trt A = 0.25, Trt C = 0.31)]. The range of population %CV values used in these simulations were described in Methods. Pairwise comparisons included Trts A (n=5 per set) vs C (n=3 per set) and Trt B vs C. For the latter sets of pairwise comparisons, all sets included n=5 randomly selected lots per treatment since there was no clearly defined reference.

#### Challenging equivalence between Trt B or C vs Trt A: PRZ

Using the mean PRZ MCI values from Table 3, the %CV for the first set of population simulations were Trt A = 20% and Trt C = 10%. With that magnitude of variability, significant differences were only detected for Set 1 (Table 6a). By decreasing the specified Trt A simulation %CV to 10, statistically significant differences were identified in all but one set (Table 6b). The set not exhibiting statistical significance would still have been declared inequivalent based upon the presence of one lot of Trt C that failed to overlap with those of Trt A.

**Table 6a:** The %CV of the randomly selected Trt A lots resulted in all but one set where Trt A %CV >16. Population simulations specified 20% CV Trt A (n=5 per set) and 10% CV Trt C (n=3 per set) and mean values of Trt A = 0.25, Trt C = 0.30.

|  | S1 Trt A | Trt C | S2 Trt A | Trt C | S3 Trt A | Trt C | S4 Trt A | Trt C | S5 Trt A | Trt C |
| --- | --- | --- | --- | --- | --- | --- | --- | --- | --- | --- |
| mean | 0.200 | 0.305 | 0.288 | 0.327 | 0.256 | 0.316 | 0.264 | 0.307 | 0.260 | 0.327 |
| %CV | 7.84 | 12.58 | 16.58 | 6.24 | 25.05 | 2.14 | 18.30 | 3.26 | 24.74 | 6.24 |
| Ratio | 1.52* |  | 1.14 |  | 1.23 |  | 1.16 |  | 1.26 |  |

\*P<0.05.

**Table 6b:** The %CV of the randomly selected Trt A lots resulted in all sets of Trt A %CV <10. Population simulations specified 10% CV Trt A (n=5 per set) and 10% CV Trt C (n=3 per set)) and mean values of Trt A = 0.25, Trt C = 0.30.

|  | S1 Trt A | Trt C | S2 Trt A | Trt C | S3 Trt A | Trt C | S4 Trt A | Trt C | S5 Trt A | Trt C |
| --- | --- | --- | --- | --- | --- | --- | --- | --- | --- | --- |
| mean | 0.255 | 0.305 | 0.254 | 0.327 | 0.252 | 0.316 | 0.253 | 0.307 | 0.252 | 0.327 |
| %CV | 9.29 | 12.58 | 6.45 | 6.24 | 6.97 | 2.14 | 9.08 | 3.26 | 8.99 | 6.24 |
| Ratio | 1.19 |  | 1.30* |  | 1.25* |  | 1.22* |  | 1.30* |  |

\*P<0.05

When Trt A was simulated at 20% CV, in addition to S1, S2 would also have failed to demonstrate comparability because of one lot of Trt C that fell slightly outside the range of Trt A values (Figure 5). When the Trt A %CV was reduced to 10%, all sets but one were significantly different. Nevertheless, that set would have failed to demonstrate equivalence because of one Trt C lot that fell outside the range values associated with Trt A (captured in Figure 5 as LoCVS1TrtA vs Trt C). Therefore, as shown in the previous challenge, the ability to identify treatment differences decreases when the reference product %CV is higher than that of the test treatment.

**Figure 5:**
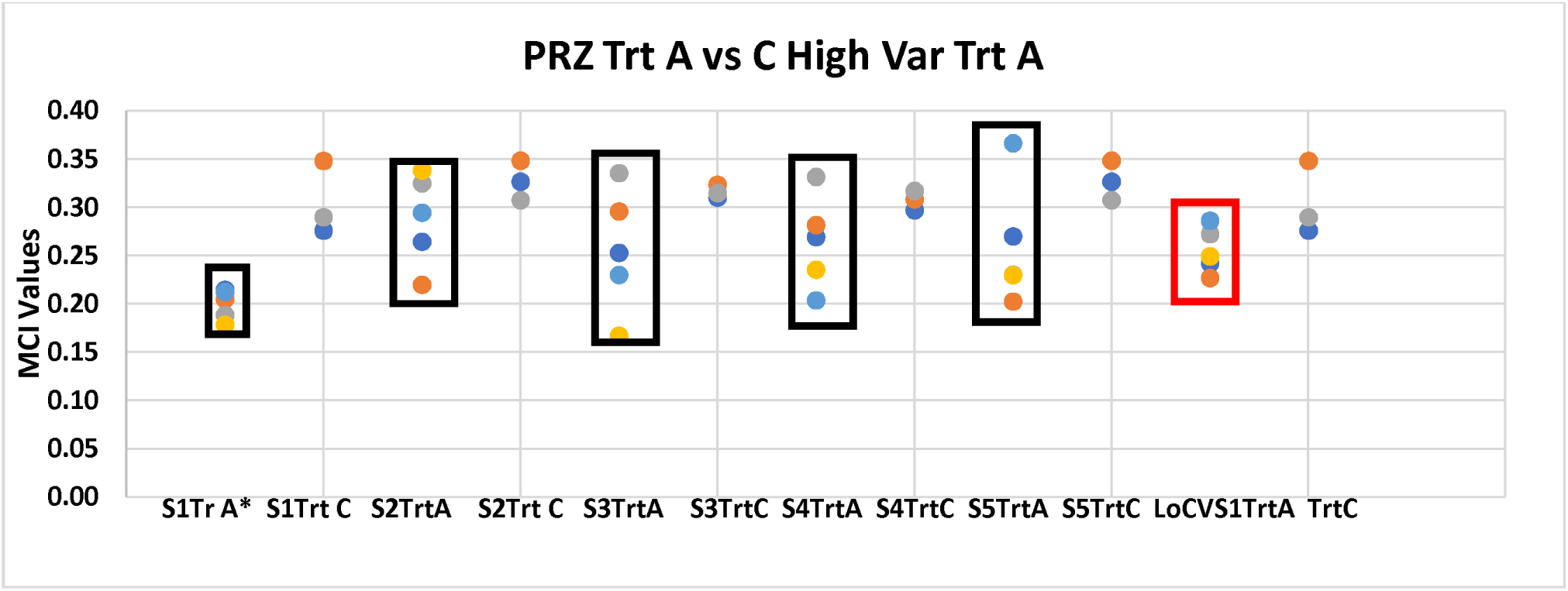
PRZ Trt A %CV = 20 (S1-S5, black box) and the one set where Trt A %CV=10 (red box, n=5 per set) vs Trt C (n=3 per set) were not declared statistically significant. Trt C %CV = 10 across all comparisons. Population mean Trt C = 0.31 and Trt A = 0.25.

#### Challenging equivalence between Trt B or C vs Trt A: IVM

Since the Trt A MCI value is the same as that for PRZ (MCI = 0.25) and while that for Trt C is slightly lower (IVM Trt C = 0.30, PRZ Trt C =0.31), a different set of %CV values were used for these simulations. In this case, the objective was to determine if a low %CV for Trt C (%CV = 1 or 10) could overcome the high %CV for Trt A (20%) and identify statistically significant differences.

Statistical significance was detected for all sets when Trt C was simulated with CV = 1%. Three out of five sets were associated with P<0.05 when the Trt C simulations were increased to CV= 10%. When plotting the individual MCI values for the two sets associated with P>0.05, at least one Trt C test value fell outside the range of Trt A MCI values (Table 7, Figure 6 Sets 2 and 5). Therefore, none of these comparisons would have been declared comparable.

**Figure 6:**
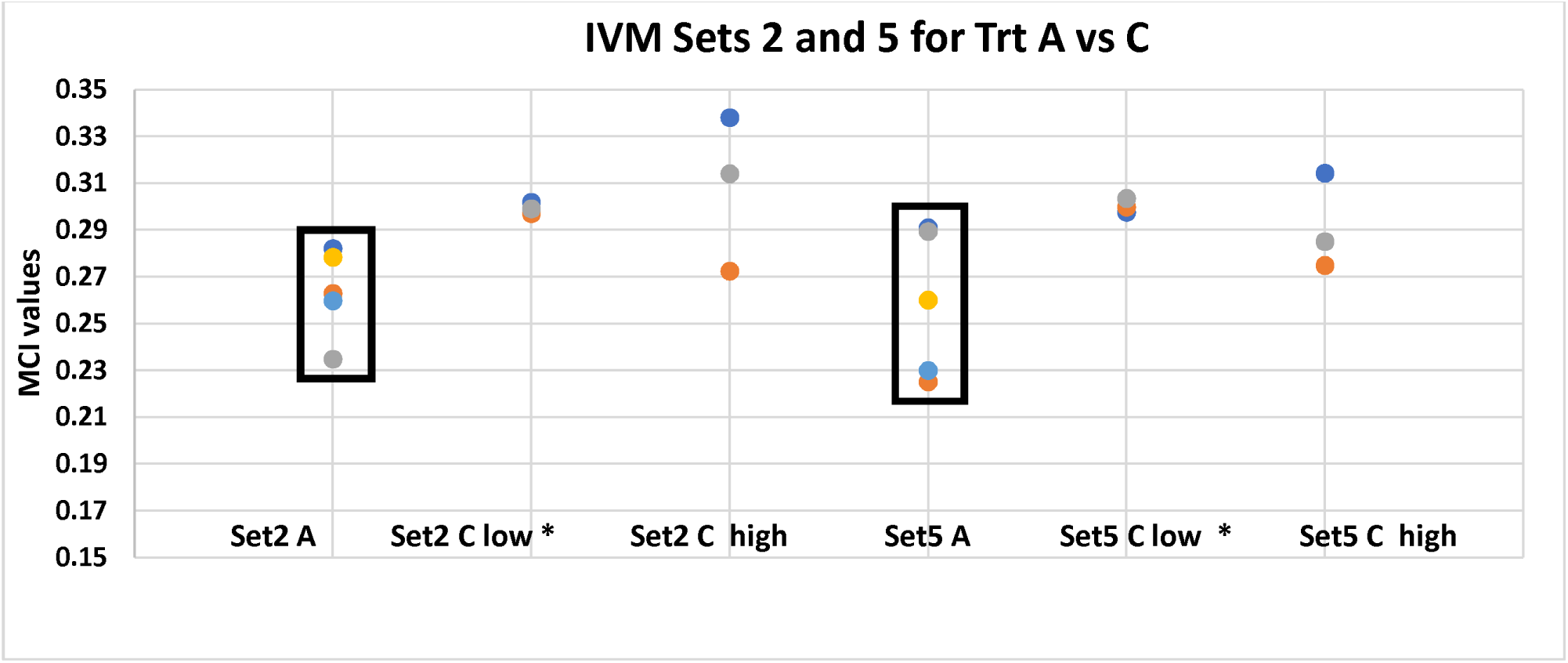
Distribution of values for sets not statistically significant when Trt C %CV=10. IVM was Trt C simulated at 1% or 10% CV and Trt A = 20% CV. Sets highlighted with black boxes represent Trt A. Population means simulated as Trt A = 0.25, Trt C = 0.30.

**Table 7:** IVM Trt A (n=5 per set) vs C (n=3 per set), where Trt C simulated at 1% or 10% CV, mean 0.30 and Trt A = 20% CV, mean 0.25.

| IVM | S1<br>Trt A | Trt C<br>1%<br>CV | Trt C<br>10%<br>CV | S2 Trt<br>A | Trt C<br>1%<br>CV | Trt C<br>10%<br>CV | S3 Trt<br>A | Trt C<br>1% CV | Trt C<br>10% CV | S4 Trt<br>A | Trt C<br>1%<br>CV | Trt C<br>10% CV | S5 Trt<br>A | Trt C<br>1% CV | Trt C<br>10% CV |
| --- | --- | --- | --- | --- | --- | --- | --- | --- | --- | --- | --- | --- | --- | --- | --- |
| mean | 0.230 | 0.301 | 0.291 | 0.264 | 0.299 | 0.308 | 0.252 | 0.301 | 0.31166 | 0.250 | 0.299 | 0.3461 | 0.259 | 0.300 | 0.291 |
| %CV | 19.18 | 0.71 | 7.64 | 7.11 | 0.81 | 10.78 | 10.14 | 1.43 | 7.02 | 5.59 | 1.04 | 9.21 | 12.11 | 0.99 | 7.03 |
| Ratio |  | 1.31* | 1.27* |  | 1.14* | 1.17 |  | 1.20* | 1.24* |  | 1.20* | 1.39* |  | 1.16* | 1.12 |

#### Challenge 2: Potential for declaring IVM or PRZ Trt B comparable to Trt C

For this evaluation the “test product” n values were increased to 5 (same as the reference) because of the larger treatment differences in MCI values.

#### Comparing Trt B and C: PRZ

The question was whether differences would have been detected in their respective MCI values since Trts B and C *in vivo* BE study results were only slightly outside the traditional confidence limits for Cmax and AUC0-2 (0.88 – 1.27) but not for AUC0-last (0.81-1.08).

The Monte Carlo simulation of population MCI values were as defined in Table 3b (Trt B = 0.33; Trt C = 31). Trt B MIC values were simulated with a 10% CV and Trt C with either 1% or 10% population CV. Statistically significant differences were detected for two of the 5 sets at Trt C population %CV = 1, and none of the sets with P≥0.05 had Trt C MCI values failing to overlap with those of Trt B. When the Trt C population %CV was increased to10, none of the of 5 sets were associated with statistically significant differences. However, two sets would not have been declared comparable (S1 and S5) due to at least one Trt C lot with MCI values outside the range of those for Trt B (Table 8 and Figure 7).

**Figure 7:**
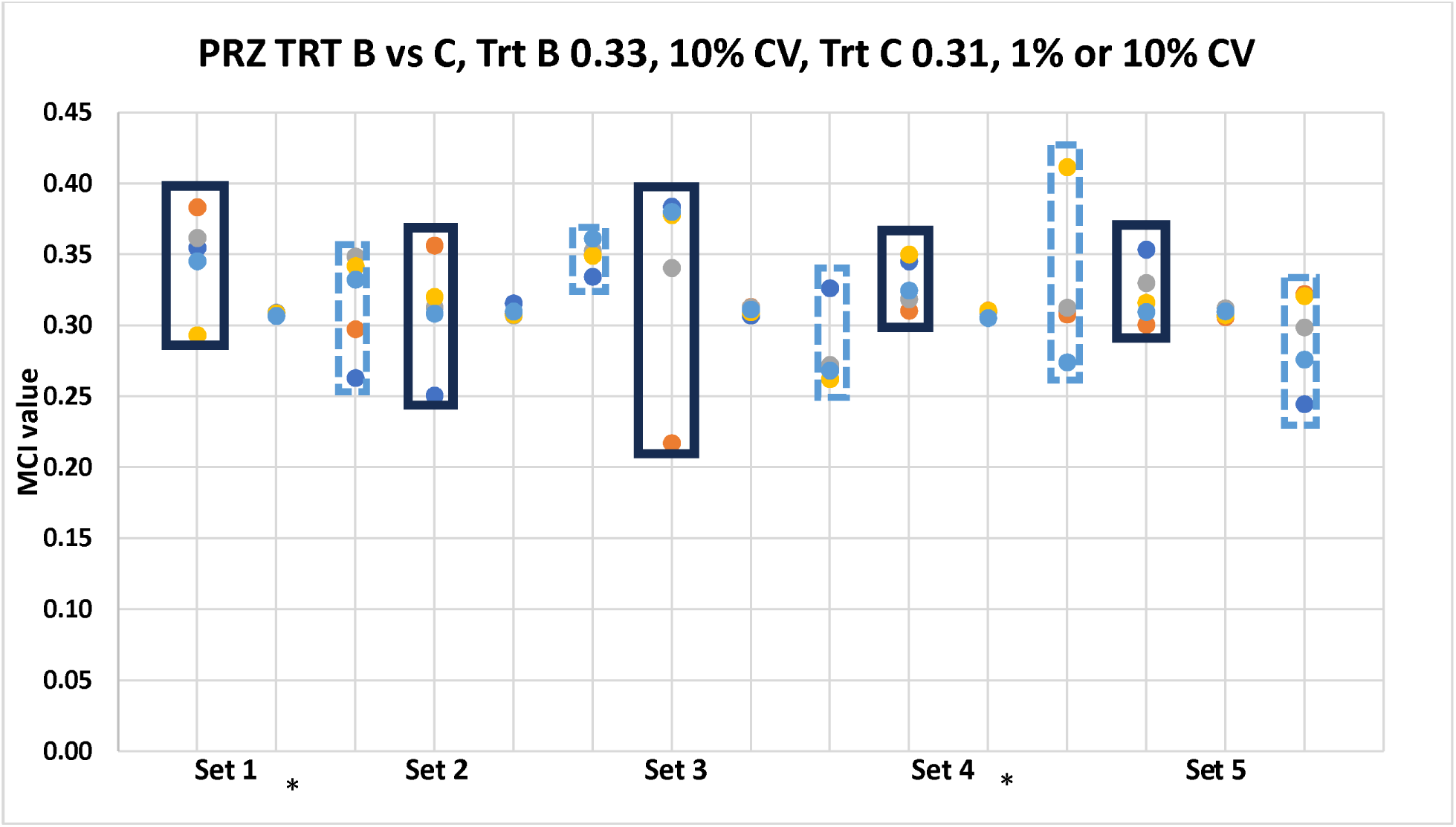
PRZ TRT B vs C, Trt B mean = 0.33, 10% CV, Trt C mean = 0.31, 1% or 10% CV. For each set, lots highlighted with black box are Trt B, no highlight is Trt C 1% and blue dashed box is Trt C 10% (for each set, n=5 per treatment). * P<0.05

**Table 8:** Means and %CV for randomly selected Trt B and C “lots” for each set of comparisons (for each set, n=5 per treatment).

| PRZ | S1TRT B | Trt C 1% | Trt C 10% | S2TRT B | TRT C 1% | Trt C 10% | S3TRT B | Trt C 1% | Trt C 10% | S4TRT B | Trt C 1% | Trt C 10% | S5TRT B | Trt C 1% | Trt 10% |
| --- | --- | --- | --- | --- | --- | --- | --- | --- | --- | --- | --- | --- | --- | --- | --- |
| Mean | 0.348 | 0.308 | 0.317 | 0.310 | 0.310 | 0.349 | 0.340 | 0.311 | 0.280 | 0.330 | 0.309 | 0.323 | 0.322 | 0.309 | 0.29 |
| %CV | 9.64 | 0.36 | 11.34 | 12.28 | 1.12 | 2.79 | 20.83 | 0.84 | 9.40 | 5.20 | 0.72 | 16.03 | 6.41 | 0.79 | 11.1 |
| Ratio Trt C/B |  | 0.89* | 0.91 |  | 1.00 | 1.13 |  | 0.91 | 0.82 |  | 0.94* | 0.98 |  | 0.96 | 0.9 |

\*P<0.05

The small margin between failing and passing was consistent with the marginal difference between Trt B and C *in vivo* Cmax and AUC0-2 values.

#### Comparing Trt B and C: IVM

The relationship between Trt B (population mean = 0.47) and C (population mean = 0.30) was examined when the simulated population %CV was 20% for Trt B and 10% for Trt C. Given the large difference between IVM Trt B and C MCI values, the large Trt B %CV was intended to challenge the likelihood of identifying statistically significant treatment differences (Table 9). All but one comparison was associated with P<0.05 (S4), indicating the ability to identify large differences in MCI values even when variability is high (Table 9 and Figure 8).

**Figure 8:**
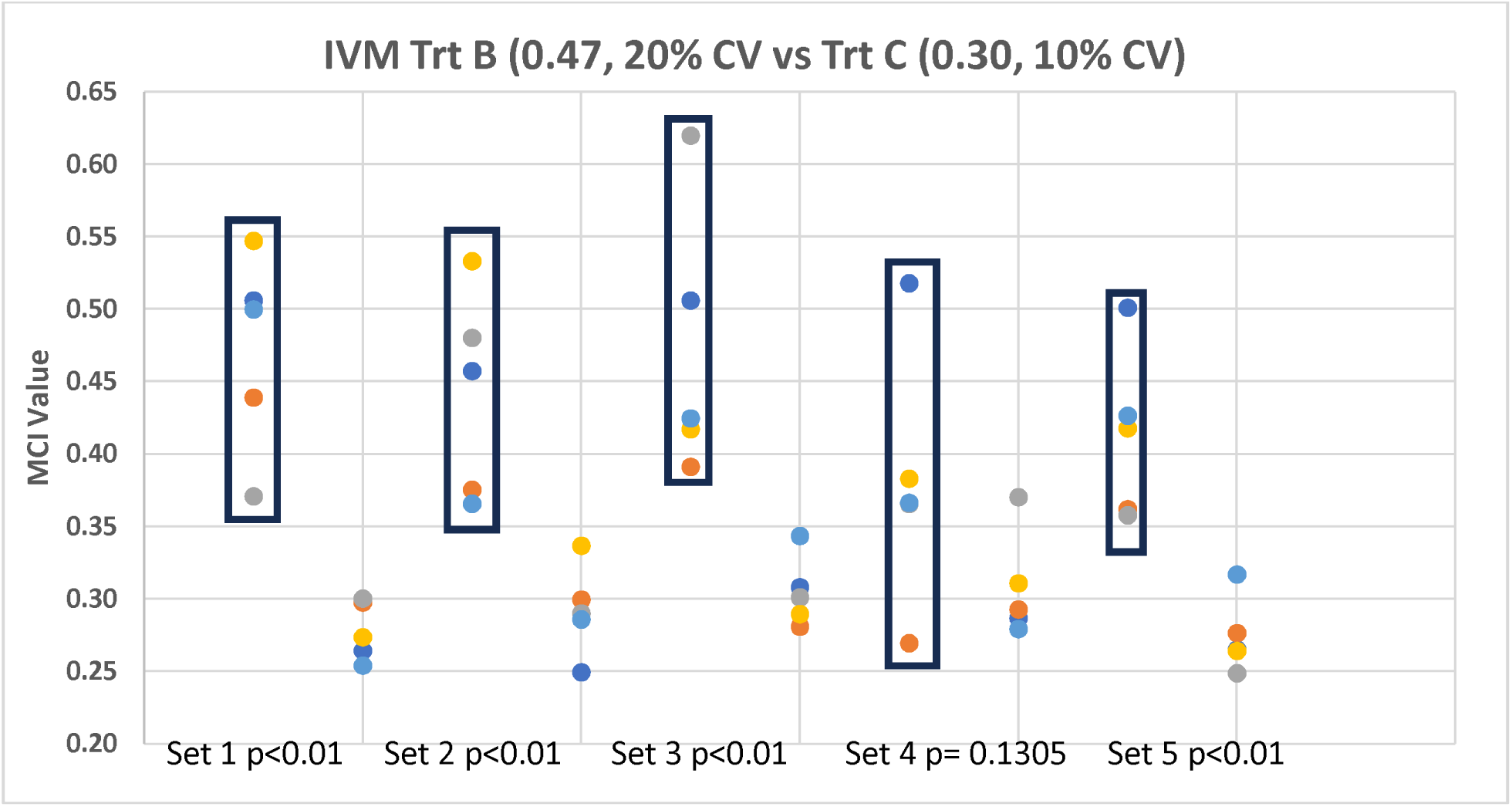
IVM Trt B (mean =0.47, 20% CV vs Trt C (mean =0.30, 10% CV) (for each set, n=5 per treatment).

**Table 9:** The relationship between TRT B (, simulated mean = 0.47, CV=20%) and C (simulated mean = 0.30, CV = 10%) (for each set, n=5 per treatment).

| IVM | S1 Trt B 20% | Trt C 10% | S2 Trt B 20% | Trt C 10% | S3 Trt B 20% | Trt C 10% | S4 Trt B 20% | Trt C 10% | S5 Trt B 20% | Trt C 10% |
| --- | --- | --- | --- | --- | --- | --- | --- | --- | --- | --- |
| Mean | 0.473 | 0.278 | 0.442 | 0.292 | 0.472 | 0.305 | 0.380 | 0.308 | 0.413 | 0.274 |
| %CV | 14.54 | 7.33 | 16.09 | 10.70 | 19.76 | 7.91 | 23.37 | 11.92 | 14.12 | 9.41 |
| Ratio C/B | 0.59* |  | 0.66* |  | 0.65* |  | 0.81 |  | 0.66* |  |

\*P<0.05

## Discussion

The utility of MoCMC system for detecting inequivalent formulations is dictated by the parameters selected for inclusion in the polygon. While it is possible that differences could be detected between treatment MCI values even if those differences lack any *in vivo* relevance, this risk could be mitigated by a careful selection of parameters that are aligned with the product CQAs. This necessitates formulation understanding and typically, the generation of pilot batches across a range of formulations and manufacturing specifications.

It is also necessary to characterize the magnitude of product Q3 interbatch variability. While high reference product variability may imply that a wider difference in treatment MCI values will not compromise product comparability, high variability would likely be problematic when associated with the test formulation. Regardless of the source, high variability will negatively influence statistical power, rendering it difficult to detect product differences. For this reason, in addition to the statistical test for differences, it was necessary to compare the ranges of test and reference MCI values. Nevertheless, the statistical gap that currently exists has the potential to detract from the utility of the MoCMC to support a conclusion of product BE for locally acting drug products. We initially considered the possibility of generating a tolerance limits approach for describing the likely range of reference MCI values (e.g., 99% of the population with 95% confidence). ^13^ However, this was quickly rejected because of the width of the interval that would be generated due to the small number of reference lots likely to be available. Therefore, we are currently exploring other alternatives for incorporating a statistical test of equivalence into the MoCMC assessment. As part of that assessment, we are also examining the manner in which the ratio of treatment MCI values (and therefore direction and width of the confidence interval) might be influenced by the ordering of the parameters defining the areas of irregular polygons.

Another observation was that the sensitivity to product differences in a particular parameter decreases as the number of radii increase. Since the MoCMC handles all attributes as equally important, if differences in a particular attribute are of concern, that parameter should be evaluated outside the MoCMC. For example, results of the pivotal dissolution comparison should be handled as a stand-alone variable.

Although the *in vitro* dissolution of Trts A and B were found to be comparable (Martinez et al., 2024), it is likely that tablet equivalence would not have been concluded if the MoCMC system was applied across multiple lots (see Table 6b and Figure 5). Conversely, there is the possibility that Trts B and C would have been declared equivalent (Table 8). Such a possibility may not negatively align with a conclusion of comparable local *in vivo* drug exposure since the *in vivo* study Trt B vs C were within the limits of 0.80 – 1.25 for AUC0-last and only slightly outside the conventional 90% confidence interval limits for Cmax and AUC0-2 (0.88-1.27).

For IVM, the results of the MoCMC evaluation were consistent with the *in vivo* BE study in that none of the pairwise comparisons succeeded in being equivalent to Trt A. More challenging was the *in vivo* similarity between Trts B and C. While the 90% confidence interval for Trts B vs C would have been within the BE criteria for AUC0-last and only slightly outside the BE criteria for Cmax (0.91-1.26), it was well outside the acceptable 90% confidence limits for AUC0-3 (1.01-1.41; this parameter was included in our *in vivo* data analysis since it could reflect product differences in the *in vivo* exposure of parasites residing within the upper GI segments). The MoCMC would have successfully caught this potential source of inequivalence.

It is worthwhile noting that the magnitude of differences in IVM MCI values exceeded those of PRZ. Nevertheless, the *in vivo* profile differences for PRZ exceeded those seen for IVM. This indicates that the magnitude of MCI differences should not be interpreted as correlating with the extent of product *in vivo* inequivalence. Rather, this tool should be limited to addressing whether, within the scope of the selected Q3 parameters, a test and reference product *in vivo* dissolution behavior are likely to be similar That the difference in MCI values between Trts B vs C and A vs B were substantially greater for IVM than they were for PRZ indicates that either: 1) the selection of Q3 parameters were not equally reflective of the CQAs for these two APIs, or 2) there was a disproportionate emphasis on *in vitro* dissolution (a function of the absence of information available for the other Q3 parameters). Without the *in vivo* and *in vitro* testing of additional formulations, we cannot further examine reasons for differences in the PRZ vs IVM MCI treatment comparisons.

Other parameters that could have been highly informative include granule size distribution within each sieve cut (to improve the assessment the average % theoretical concentration for each drug within each treatment), hardness, and friability. It would also have been informative to have had information on the size of the granules after tablet compaction. One of the possibilities hypothesized after using a modeling and simulation approach to explore factors that may have contributed to the observed inequivalence of Trt A was that of granule agglomeration when the smallest (sticky) particles were exposed to the tableting compaction force. The impact of that agglomeration may not have been evident during in the *in vitro* dissolution study due to the volume of media used relative to that in the fasted dog stomach and to the agitation-associated hydrodynamics present inside the vessels when using the USP paddle at 50 rpm. This possibility could have been explored if the tablets were still available for testing using a non-destructive method (e.g., UV/IR absorption)^14,15^.

## Conclusion

This manuscript sets a foundation for identifying factors that can potentially influence the results obtained when incorporating the MoCMC approach as part of a product in vitro BE assessment. The ultimate goal is that with this understanding, we can make the adjustments necessary for it to serve as one of the pivotal components in a totality of evidence approach for evaluating the BE of products containing nonsystemically absorbed drugs. When combined with formulation (Q1 and Q2 sameness) and the pivotal *in vitro* dissolution comparison, an in vitro approach for the determination of product BE may be considered in lieu of clinical endpoint BE studies for orally administered, non-systemically absorbed drug products.

## Acknowledgement

The authors wish to express their appreciation to Dr. Xin Fang, Supervisory Statistician, CVM, for his statistical advice.

## Disclaimer

The findings and conclusions in this research have not been formally disseminated by the Food and Drug Administration and should not be construed to represent any Agency endorsement, determination, or policy. The mention of commercial products, their sources, or their use in connection with material reported herein is not to be construed as either an actual or implied endorsement of such products by the FDA.

## Author Contributions

Marilyn Martinez and David Longstaff shared in the data analysis and manuscript development.

## Funding Formulation

No funds were received in support of this work.

## Data Availability

The in vivo and in vitro datasets are available upon request.

## SUPPLEMENTAL INFORMATION

Attributes used in the MoCMC evaluation of the PRZ/IVM treatment comparisons:

A. Dissolution:
  1. The % dissolved at 20 minutes. The 20-minute sampling time was selected because release was still increasing rapidly at that timepoint. Given that the dissolution in phosphate buffer was considered the pivotal method, those data were analyzed for F2 metric outside of the MoCMC to be used as a confirmatory test. This left 2 other methods for inclusion in the MoCMC:
    a. 0.1 N HCl + 0.2% SLS
    b. pH 4.5 acetate buffer, 0.5% SLS
  2. The amount of PRZ dissolved at the last dissolution timepoint (120 minutes).
    a. 0.1 N HCl + 0.2% Sodium Lauryl Sulfate (SLS)
    b. pH 4.5 acetate buffer, 0.5% SLS
  3. For IVM, dissolution in 0.1 N HCL + 0.2% SLS, the % dissolved reached a maximum amount and subsequently declined for Treatments A and C. However, this did not occur with Treatment B where the %dissolution continued to increase throughout the 120-minute duration of sampling. This decline did not occur in acetate buffer. Therefore, the parameters included for IVM were as follows:
    a. % dissolved at 120 min/maximum amount dissolved in 0.1 N HCl + 0.2% SLS
    b. % dissolved in pH 4.5 acetate buffer, 0.5% SLS at 120 minutes.
B. Granule information:
  4. For each drug, the 2 parameters were:
    a. granule size
    b. % theoretical concentration within each granule were included in the assessment.

### In Vitro Dissolution

The pivotal *in vitro* dissolution tests were conducted in 500 mL buffer, paddle 50 RPM, in the following media:

0.1N HCl + 0.2% SLS (n = 6 per formulation)
0.1 M acetate buffer pH 4.6 + 0.5% SLS (n = 6 per formulation)
0.01 M phosphate buffer, pH 6.8 + 0.5% SLS (n = 12 per formulation). The data collected in phosphate buffer were not included in the MoCMC since they were analyzed separately for calculating the F2.

Release of the APIs in each vessel was determined from a single aliquot of the medium at 10, 20, 30 60, 90, and 120 minutes (min). The analytical method simultaneously quantified PRZ and IVM using a gradient UPLC system with detection at 245 nm (Hollenbeck et al., 2025).

### Determining the Granule Particle Size

The method by which the three tablet formulations were manufactured and the corresponding composition of each tablet was defined by Hollenbeck et al, 2025. Part of the distinction between the modified release and the fast release tablets was the size of the granules and the corresponding % theoretical concentration contained within each granule.

Each granulation was classified into the following sieve cuts: 18/20 (840–1,000 μm), 20/40 (420–840 μm), and 40/60 (250–420 μm). Given the inconsistent relationship between granule size and % theoretical concentration of PRZ or IVM as a function of granule, it was necessary to determine how best to translate this attribute to a radius for each formulation. The tablets manufactured using a combination of the small particle size granules of the immediate release formulations [i.e., praziquantel fast release = PF (sieve cut 40/60) and ivermectin fast release = IF (sieve cut 40/60)] was chosen as a reference formulation (Trt A). The other sieve cuts were PF (20/40) and ivermectin medium release = IM (20/40) for the manufacture Trt B, and the praziquantel medium release = PM (sieve cut 18/20) and IF (sieve cut 20/40) for the manufacture of Trt C.

Hollenbeck et al., 2025 also determined the % theoretical drug concentration within the granules as a function of mean particle size (Table 1). These values can be used to determine the calculate the tablet label claim rather than simply use values in the table. For example, if we used about 300 μm for the fast PRZ, you could see that the value will be markedly than PM. Similar point for the IVM.

Hollenbeck et al. reported that the granule size did not show a consistent relationship with the % theoretical concentration of either PRZ or IVM. Therefore, in the absence of data describing the distribution of granule sizes within each sieve cut, we needed to assume a comparatively equal distribution within a given sieve cut. Accordingly, the granule diameter used in our assessment was calculated as the middle value (0.5* largest + smallest particle size) for a given sieve cut. The corresponding % theoretical concentration was calculated as the average % theoretical concentration across the range of granule sizes associated for a specified sieve cut. For example, for the PF granules with sieve cut 40/60, our estimated % theoretical concentration was that determined by Hollenbeck et al. for mean particle size (µm) of 275, 325, 375, and 450 (Table 1).

**Supplemental Table 1:** Determination of the radii values for the test formulations.

| Parameter | Ref | Test |
| --- | --- | --- |
| % Dissolved at 20 min in HCl | 5 | (Test at 20 min/Ref at 20 min)$\times 5$ |
| % Dissolved at 20 min in Acetate Buffer | 5 | (Test at 20 min/Ref at 20 min)$\times 5$ |
| Granule Size | 5 | (Test /Ref )$\times 5$ |
| % Theoretical concentration of the granule | 5 | (Test /Ref )$\times 5$ |
| % Dissolved at 120 min in HCl | 5 | (Test at 120 min/Ref at 120 min)$\times 5$ |
| % Dissolved at 120 min in Acetate Buffer | 5 | (Test at 120 min/Ref at 120 min)$\times 5$ |

**Table 2:**
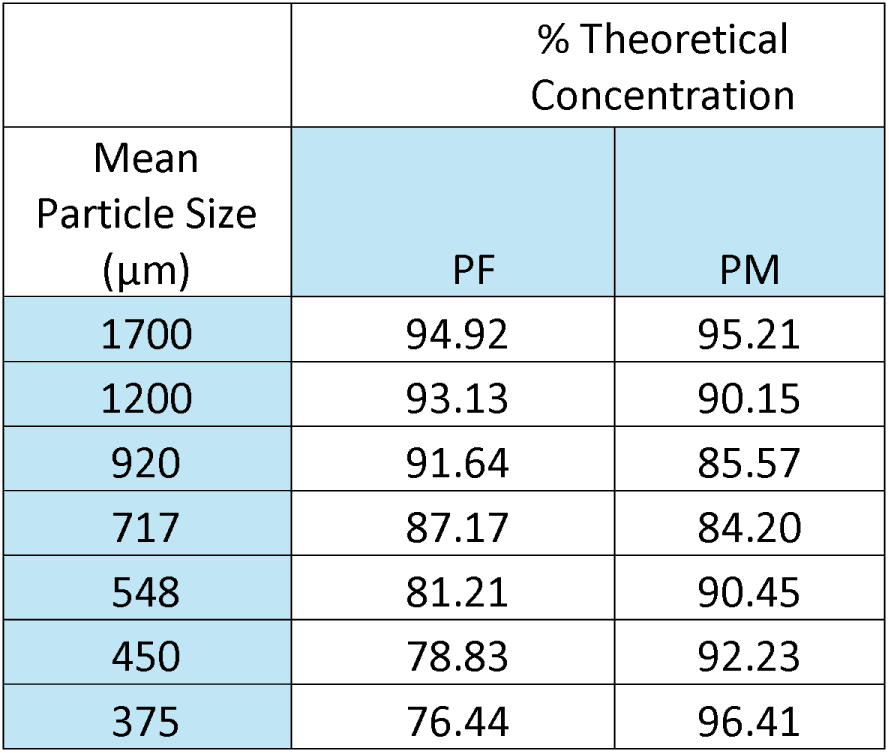
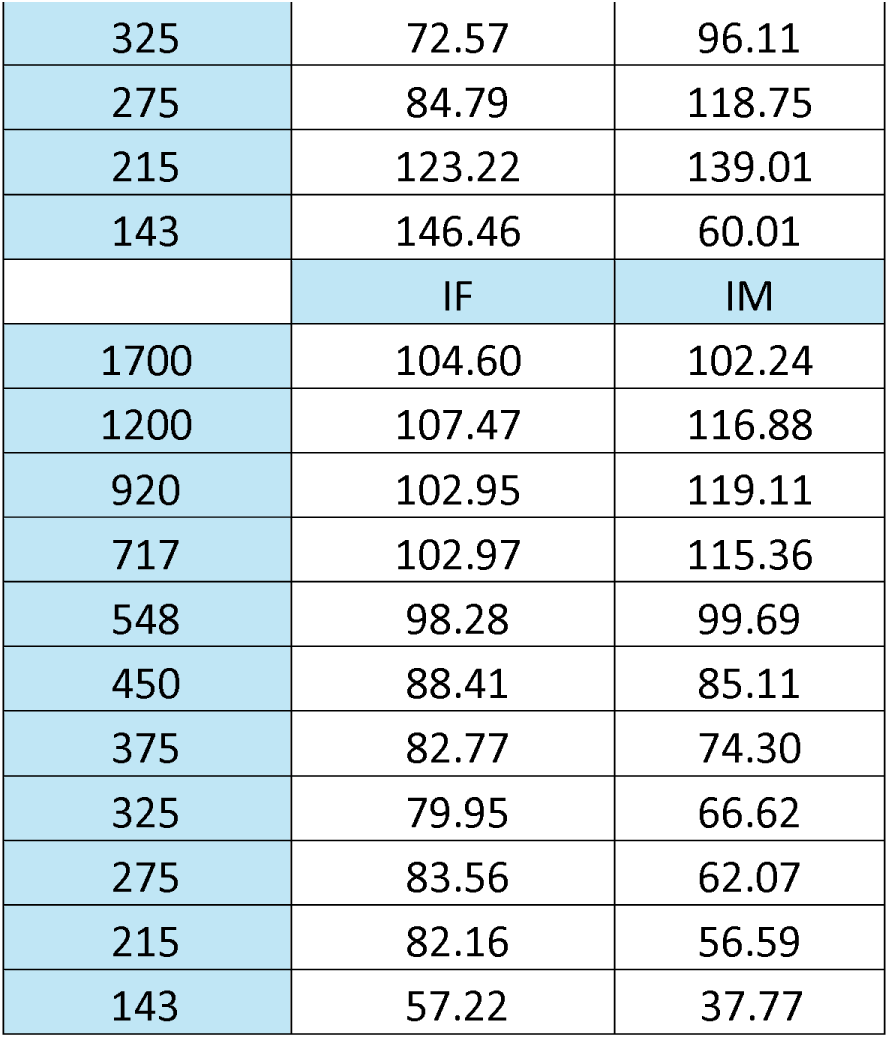
% theoretical concentration as a function of granule size and drug.

